# Environmental change effects on life history traits and population dynamics of anadromous fishes

**DOI:** 10.1101/577262

**Authors:** P. Catalina Chaparro-Pedraza, André M. de Roos

## Abstract

Migration, the recurring movement of individuals between a breeding and a non-breeding habitat, is a widespread phenomenon in the animal kingdom. Since the life cycle of migratory species involves two habitats, they are particularly vulnerable to environmental change, which may affect either of these habitats as well as the travel between them. In this study, we investigate the consequences of environmental change affecting older life history stages for the population dynamics and the individual life history of a migratory population. In particular, we use a theoretical approach to study how increased energetic cost of the breeding travel and reduced survival and food availability in the non-breeding habitat affect an anadromous fish population. These unfavorable conditions have impacts at individual and population level. First, when conditions deteriorate individuals in the breeding habitat have a higher growth rate as a consequence of reductions in spawning that reduce competition. Second, population abundance decreases, and its dynamics change from stable to oscillations with a period of four years. The oscillations are caused by the density-dependent feedback between individuals within a cohort through the food abundance in the breeding habitat, which results in alternation of a strong and a weak cohort. Our results explain how environmental change, by affecting older life history stages, has multiple consequences for other life stages and for the entire population. We discuss these results in the context of empirical data and highlight the need for mechanistic understanding of the interactions between life history and population dynamics in response to environmental change.

## Introduction

Animals from all major animal taxa set out every year on a journey between habitats in step with seasonal changes. Generally, these seasonal changes also define the timing of breeding; therefore, the habitats occupied by animals during different seasons are known as breeding and non-breeding habitat. Since the two habitats frequently differ as much as the freshwater and the marine habitat, migratory individuals are subject to a broad range of environmental influences. Exposure to such influences makes them particularly vulnerable to environmental change affecting the habitats they use. Furthermore, given the spatial separation between the habitats and the wide variety of conditions among them, the effect of environmental change differs across the habitats used by individuals at different stages during their life cycle. In consequence, a change in ecological conditions in one of the habitats directly impacts individual life history. Life history, in turn, has large effects on population processes (de Roos & Persson, 2013), which strongly influence the realized life history of organisms through the population feedback on, for example, food conditions (de Roos, Persson, & McCauley, 2003). Therefore, environmental change has the potential to affect the interaction between life history and population ecology of migratory, and in particular, anadromous species.

Anadromous individuals begin their life cycle in the breeding habitat in freshwater and later migrate to the non-breeding habitat in the ocean (this migration we from here on refer to as habitat switch), where they grow larger and eventually become mature. Mature individuals migrate back to the breeding habitat in order to reproduce, before returning to the non-breeding habitat (hereafter breeding travel). Although anadromous fish comprise less than 1% of world fish fauna, their share in global fishery trade currently exceeds 17% and continues to grow (FAO, 2016). Many anadromous fishes including salmons, sturgeons, and shads are of both economic and cultural importance. However, like many other commercially exploited fish, they are in decline and their conservation is a major concern (Pauly et al., 2002).

Anadromous species have shown major declines in the last decades due to multiple threats such as damming, overfishing and climate change (Limburg & Waldman, 2009). Damming increases the cost of the breeding travel to the spawning grounds that negatively affects individual fecundity as it leads to larger depletion of energy reserves (Jonsson, Jonsson, & Hansen, 1997). Overfishing increases mortality in later life stage as large individuals are fishing target. And climate change, among many other impacts, is predicted to reduce ocean productivity (Hoegh-Guldberg & Bruno, 2010) which results in reduced growth rate in the marine phase of the life cycle. These threats have negative impacts on the life history of individuals by affecting their fecundity, survival and growth rate during the life stage after the habitat switch. This raises the question how these threats, by influencing the life history of individuals in a late life history stage, affect the dynamics of the entire population and, in turn, the realized life history of individuals. To address this question, we use a theoretical approach to study the consequences of increased energetic cost of the breeding travel and reduced survival and food availability in the non-breeding habitat on an anadromous population. Since these threats have an effect on the population by affecting the life history of individuals after their first habitat switch, the representation of the individual life history in our model is crucial. Hence, we use a dynamic energy budget model to describe how individuals acquire and utilize energy (Nisbet, Muller, Lika, & Kooijman, 2000) and to translate these individual-level energetics into different life history trajectories. In the present study, we use data from Atlantic salmon to parameterize the dynamic energy budget model and consider its ecology to build the model at the population level. In addition, available data of wild populations of Atlantic salmon is used to test predictions of the model.

## Methods

### The model

We formulate a model that accounts for an anadromous population migrating between two habitats (Fig. 1). We assume that in each habitat (breeding and non-breeding) the individuals exploit a different resource. The migratory population is structured by age, individual body size and energy reserves and follows semi-discrete dynamics: continuous dynamics describe the resource consumption, somatic growth, stored energy reserves and survival and a discrete map describes the pulse-wise reproduction (Persson, Leonardsson, de Roos, Gyllenberg, & Christensen, 1998).

**Figure 1.**
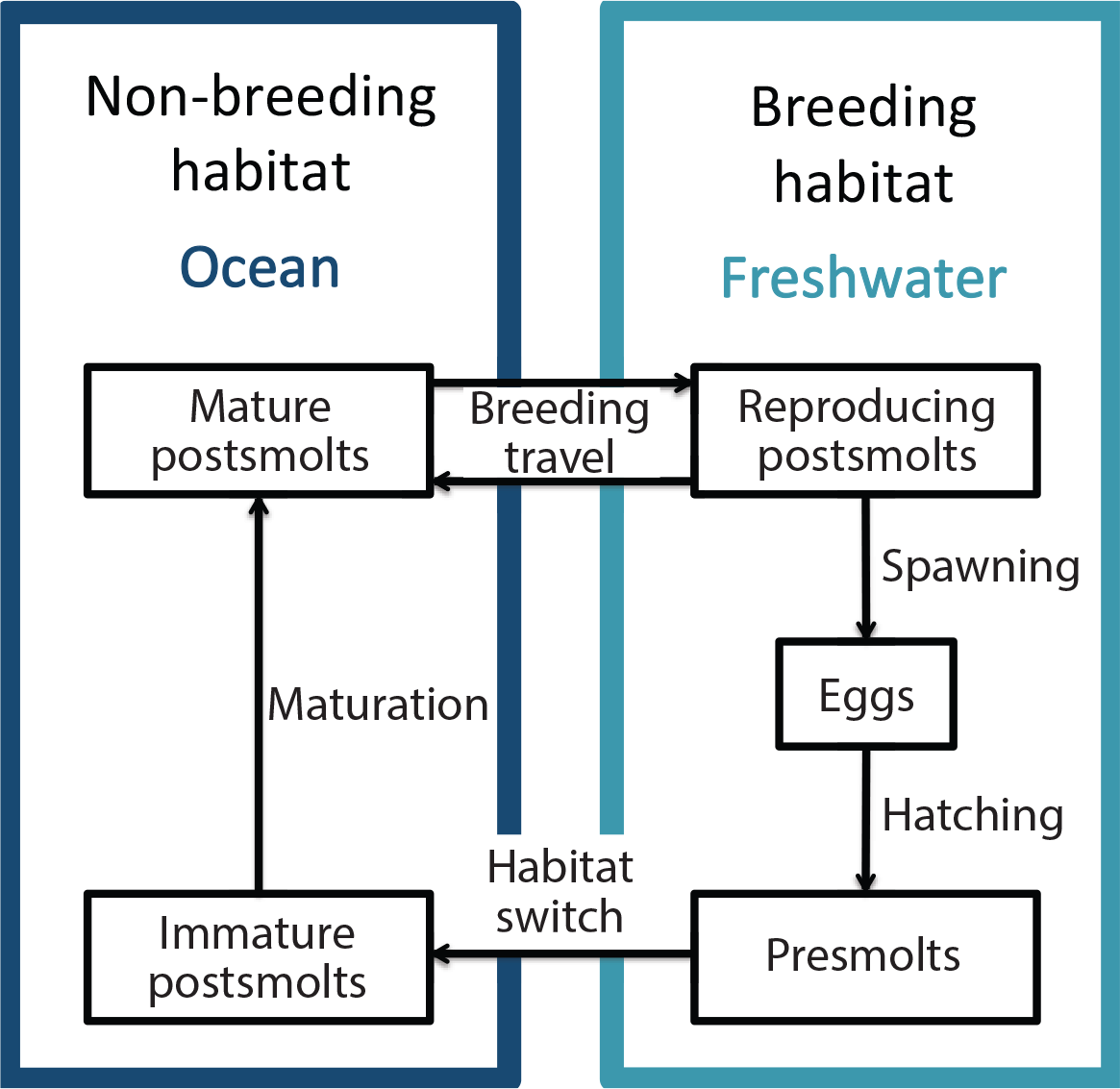
Schematic representation of the modeled anadromous life cycle

#### Yearly cycle and life history events

Salmon utilizes freshwater streams to breed, therefore its first life stages occur in this habitat. In the model female salmon are assumed to spawn in autumn at day *t*_*r*_ of the year, the eggs develop throughout winter and hatch at age *a*_*h*_ (in spring) (Hendry & Cragg-Hine, 2003). After hatching, individuals remain in the stream until they reach an age *a*_*s*_, when they smolt and migrate to the ocean (Hendry & Cragg-Hine, 2003) (hereafter, individuals younger than *a*_*s*_ are referred to as presmolts, in contrast to older individuals that are referred to as postsmolts). Every autumn, all sexually mature individuals in the ocean return to the stream and start migrating upstream at day *t*_*um*_ to spawn somewhat later at day *t*_*r*_. After spawning, postsmolts return to the ocean and finish downstream migration at day *t*_*dm*_ of the year.

#### Habitats

Density-dependence due to competition for food is strong in the breeding habitat (Jonsson, Jonsson, & Hansen, 1998), so we assumed the food density (biomass of assimilates density) to be depleted by the consumption of individuals. In the absence of consumers, the resource is assumed to follow a semi-chemostat growth dynamics with maximum density *R*_*max*_ and growth rate *ρ* (for an explanation and justification of this type of growth dynamics, see Persson et al. (1998)). Dynamics of the resource density *R*_*r*_ in the breeding habitat is hence given by:

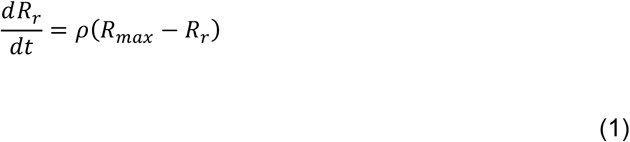

In contrast, in the non-breeding habitat, postsmolts do not experience density-dependence (Jonsson, Jonsson, & Hansen, 1998), therefore we assume a constant feeding level.

In both habitats, temperature *T* is assumed to oscillate during the year around the average temperature *T*_*m*_ with an amplitude *T*_*a*_ and a period equal to the length of the year. Therefore, the temperature reaches its maximum in summer (middle of the year) and its minimum in winter.

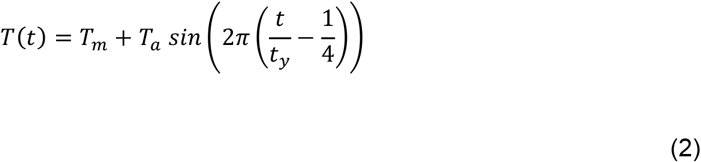

Where the *t* is the current time in days and *t*_*y*_ is the number of days of the year.

#### Individual dynamics

The core part of the model is the description of the individual behavior, that is, feeding, growth, reproduction and mortality as a function of the individual state (age, body size and energy reserves) and the state of the environment (food availability and temperature). In the following sections we describe the individual level dynamics.

##### • Feeding

In the breeding habitat, individuals are assumed to feed on the resource following a Holling type II functional response. So their feeding level *f*_*r*_ (or scaled functional response) is given by:

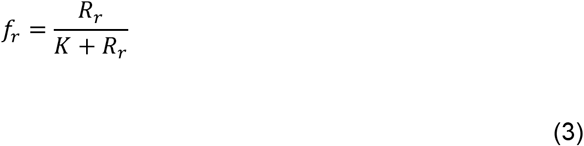

where *K* is the half saturation resource density.

While, in the non-breeding habitat density-dependence is assumed to be negligible so individuals feed at a constant feeding level *f*_*s*_.

##### • Dynamic energy budget model: Individual states

The model follows the bioenergetics approach introduced by Kooijman and colleagues (Kooijman & Metz, 1983; Nisbet et al., 2000) in which the energy allocation to somatic and reproductive metabolism are proportional to a fraction *κ* and a *1−κ* of the total energy assimilation rate, respectively. More specifically, we adopt the model developed and described in detail by (Martin, Heintz, Danner, & Nisbet, 2017). Below we provide only a concise synopsis of the model.

Individuals are characterized by three state variables: individual age *A*, structural mass *W* and energy reserves storage *S*. The acquisition and utilization of energy are described by equations (4) to (12).

The energy assimilation flux is given by:

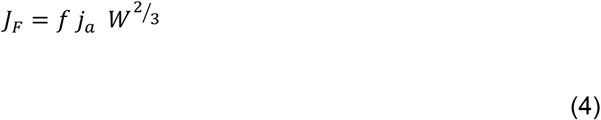

where *f* is the feeding level in either the breeding or the non-breeding habitat introduced above, *j*_*a*_ is the maximum area-specific assimilation rate and the surface area for assimilation is assumed to scale with structural mass to the power of 2/3.

Metabolic maintenance cost is the product of the mass-specific maintenance cost *j*_*m*_ and the structural mass:

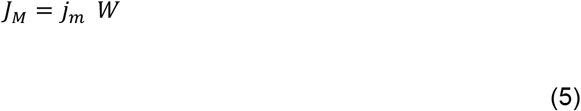

Assimilates are assumed to split into two energy fluxes: the *κ* flux and the 1 − *κ* flux. The *κ* flux is first used to cover metabolic maintenance cost, while the remaining flux *J*_*W*_ is used to synthesize structural mass. On the other hand, the 1 − *κ* flux *J*_*S*_ is allocated to energy reserves storage. Thus,

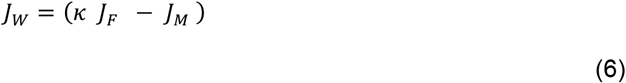

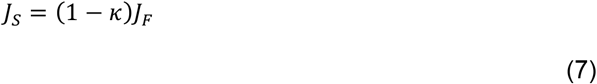

If the metabolic maintenance cost is larger than the *κ* flux, the individual starves, stops growing and depletes its storage if necessary to cover the deficit in maintenance requirements, thus in case of starvation (*J*_*M*_ > *κ J*_*F*_):

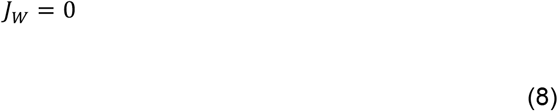

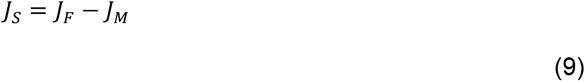

Therefore, the individual state dynamics of structural mass *W* and energy reserves storage *S* are described by the following system of differential equations:

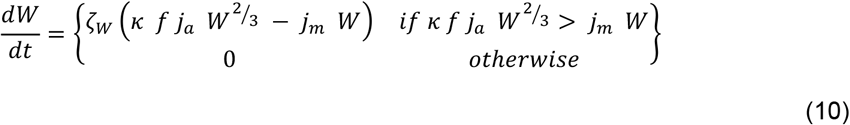

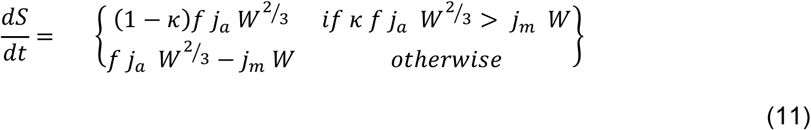

The parameter *ζ*_*W*_ in equation (10) represents the efficiency with which assimilates are converted into structural mass.

The rate constants (*j*_*a*_, *j*_*m*_) are assumed to be temperature-dependent and scale from the reference temperature *T*^*^ to the actual temperature at time *t*, *T*(*t*), following the Arrhenius relationship. Hence, both rate constants are multiplied by the temperature-correction factor *F*_*T*_(*t*):

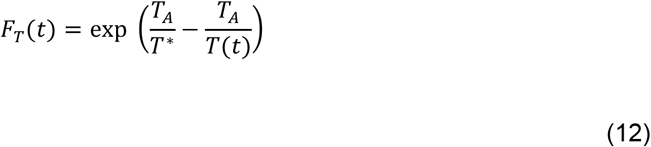

where *T*_*A*_ is the Arrhenius temperature.

##### • Maturation, reproduction and breeding travel

Individuals mature when they have reached a fixed structural mass *W*_*p*_. The storage:structure ratio at maturity *S*_*p*_/*W*_*p*_ is the threshold for reproductive investment; therefore, the excess of energy reserves above the amount that equals the *S*_*p*_/*W*_*p*_ storage:structure ratio is used for reproduction. Reproduction occurs at a discrete time *t*_*r*_. The number of offspring *θ* produced by an adult individual with structural mass *W* and energy reserves storage *S*, hence, equals:

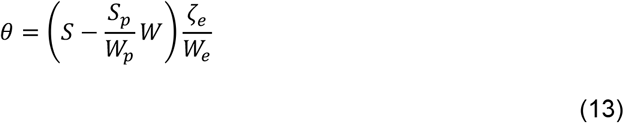

Simultaneously, the energy reserves storage value of these reproducing individuals is reduced to *S* = (*S*_*p*_/*W*_*p*_)*W*. The number of offspring produced is dependent on the yield for the conversion of storage into eggs *ζ*_*e*_ and the egg mass *W*_*e*_. During the egg stage individuals do not feed, therefore we assumed newly hatched individuals to be born with a structural mass equal to *κ W*_*b*_ and a energy reserve storage (1 − *κ*) *W*_*b*_.

During the breeding travel individuals are assumed to cease feeding, stop growing and use their energy reserves (Jonsson et al., 1997) to meet their basic metabolic maintenance cost as well as the energetic cost of the travel, which we assumed to be proportional to the metabolic maintenance cost:

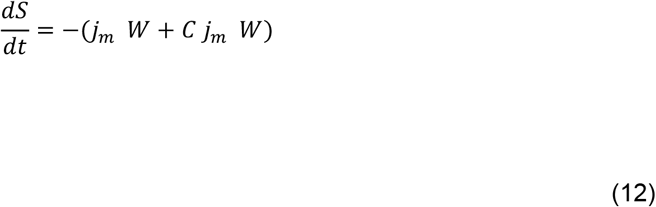

Here *C* is the proportionality constant relating the cost of the breeding travel to the metabolic maintenance cost.

##### • Survival

Individuals may die from either starvation or background mortality. The mortality rate of starving individuals with a storage:structural mass ratio *S*/*W* smaller than a threshold level q_s_ increases with decreasing values of *S*/*W*:

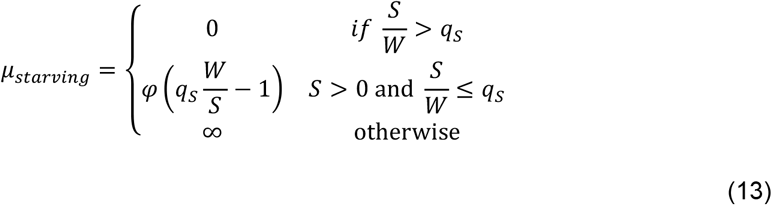

where φ is a positive proportionality constant (Persson et al., 1998). Once they have depleted their storage (S = 0) completely, starving individuals die instantaneously.

In addition to starvation mortality, individuals die at a rate *μ*_*e*_ during the egg stage, at a rate *μ*_*r*_ if they are presmolts and *μ*_*s*_ if they are postsmolts. The total per capita death rate is the sum of the different sources of mortality.

#### Population dynamics

Since reproduction occurs as a discrete event at a specific time in the year, all individuals that are born in the same reproductive event are lumped into a single cohort and assumed to grow at the same rate. Thus, we can describe the dynamics of every cohort by using a system of ordinary differential equations, which keeps track of the density of individuals, their structural mass and their energy reserves storage (see SuppInfo 1). Therefore, the dynamics of the population can be followed by numerically integrating the ordinary differential equations for each cohort separately. When a reproductive event occurs, a new cohort is added to the population, which implies additional differential equations describing the population dynamics. In addition, food density in the breeding habitat increases by an intrinsic growth process (following semi-chemostat dynamics, see Persson et al. (1998)) and decreases by consumption; these changes in food density can be followed by numerical integration of the ordinary differential equation that accounts for resource growth and consumption. The numerical integration is carried out using the EBT (Escalator Boxcar Train) (de Roos, 1988), a numerical integration method specifically designed to handle the system of differential equations that describes a physiologically structured population.

We parameterized the model for Atlantic salmon based on literature data of its individual life history and the characteristics of the breeding and non-breeding habitats. All parameter values and their sources are presented in Table 1, while SuppInfo 1 provides details about the population-level formulation of the model.

**Table 1.**
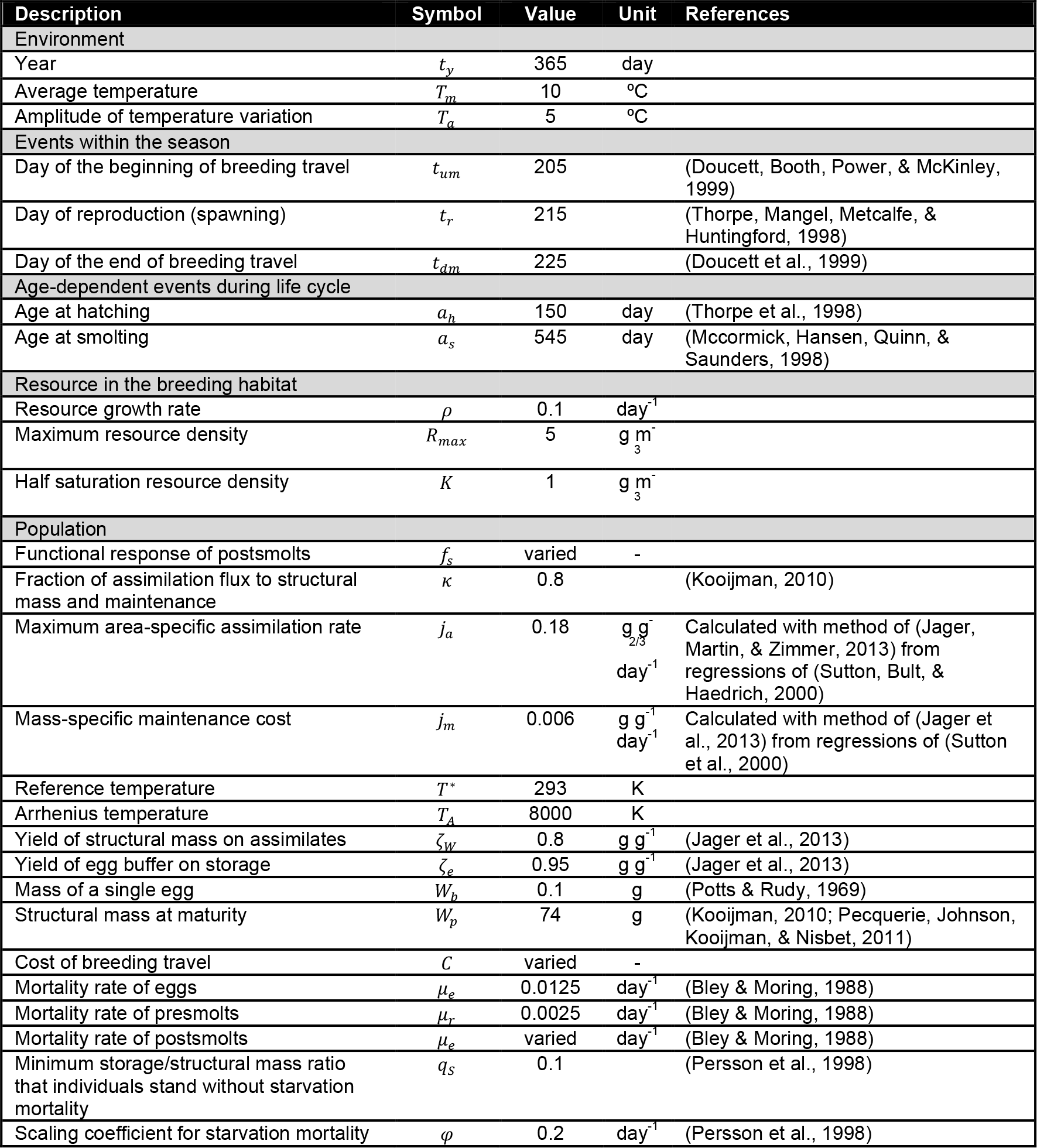
Parameter values

Our main interest is to investigate the effects of increased cost of the breeding travel and low survival and food availability in the non-breeding habitat on the population and the realized life history of individuals. To do so, we vary the feeding rate in the non-breeding habitat *f*_*s*_ between *ad libitum* food and 0.3 times the amount of food *ad libitum*, the background mortality rate of postsmolt individuals *μ*_*s*_ between 0.0063 and 0.0107 per day, equivalent to annual survival probabilities of 0.1 and 0.02, respectively, and lastly, the cost of the breeding travel *C* between 0 and 1 times the metabolic maintenance cost for the period that the breeding travel lasts (between *t*_*um*_ and *t*_*dm*_).

All model results presented correspond to the values of the population statistics after transient dynamics have disappeared.

### Data analysis

In order to test predictions of the model regarding the relationship between growth rates of presmolts and postsmolts we estimated growth rates of presmolts and postsmolts based on data of wild population of Atlantic salmon presented by Hutchings and Jones (1998) (see Table 2). The growth rate of presmolts is estimated as the difference in length between 1+ parr and smolt divided by the smolt age. Growth rate of postsmolts is estimated as the difference in length between smolts and grilse (individuals that spend one winter in the sea after smolting to migrate upstream to spawn). Hutchings and Jones (1998) present data for smolt age and length of 1+ parr, smolt and grilse for 24 populations. We tested the correlation between the growth rate of presmolts and the growth rate of postsmolts with a Pearson correlation test.

**Table 2.** Atlantic salmon life history data

| Region | River | Parr length +1 | Smolt length | Smolt age | Grilse length | Smolt-grilse survival | References |
| --- | --- | --- | --- | --- | --- | --- | --- |
| Labrador | Sandhill |  | 16 | 4.44 |  | 0.0815 | Hutchings and Jones (1998) |
| Maritimes | Miramichi | 9.39 | 12.8 | 2.8 | 55.3 |  | Hutchings and Jones (1998) |
| Newfoundland | Conne |  | 14.8 | 3.28 |  | 0.0578 | Hutchings and Jones (1998) |
| Newfoundland | Highlands | 8.27 | 13.1 | 2.9 | 53.3 |  | Hutchings and Jones (1998) |
| Newfoundland | Little Codroy | 8.59 | 15 | 2.66 | 54.1 | 0.0229 | Bley and Moring (1988), Hutchings and Jones (1998) |
| Newfoundland | NE Trepassey |  |  | 3.67 |  | 0.0622 | Hutchings and Jones (1998) |
| Newfoundland | North Hr | 8.43 | 15.1 | 3.2 | 55.1 |  | Hutchings and Jones (1998) |
| Newfoundland | West Arm Bk | 7.54 | 16.8 | 3.68 | 52.9 | 0.0489 | Hutchings and Jones (1998) |
| Newfoundland | Wings Bk | 7.63 | 16.4 | 4 | 51 |  | Hutchings and Jones (1998) |
| Quebec | Bec-Scie |  |  | 2.97 |  | 0.0283 | Hutchings and Jones (1998) |
| Quebec | Matamek | 8.59 | 15 | 3.11 | 52.9 |  | Hutchings and Jones (1998) |
| Quebec | Moisie | 6.31 | 12 | 3.57 | 48.7 |  | Hutchings and Jones (1998) |
| Quebec | Pigou | 9.62 | 15.1 | 2.93 | 53.1 |  | Hutchings and Jones (1998) |
| Quebec | Saint-Jean | 5.42 | 11.5 | 3.52 | 54.4 |  | Hutchings and Jones (1998) |
| Quebec | Trinité | 8.06 | 12.1 | 3 | 54.2 | 0.0237 | Hutchings and Jones (1998) |
| Norway | Figgenelva | 8.4 | 11.7 | 2.28 | 54.3 |  | Hutchings and Jones (1998) |
| Norway | Haelva | 8.9 | 12.5 | 2.47 | 60.2 |  | Hutchings and Jones (1998) |
| Norway | Imsa |  | 15.4 | 1.95 |  | 0.0357 | Hutchings and Jones (1998) |
| Norway | Laerdalselva | 8.2 | 11.9 | 2.82 | 67.96 |  | Hutchings and Jones (1998) |
| Norway | Mandalselva | 8.5 | 13.3 | 2.82 | 54 |  | Hutchings and Jones (1998) |
| Norway | Namsen | 8 | 11.9 | 2.67 | 58.49 |  | Hutchings and Jones (1998) |
| Norway | Nummendals | 7.2 | 10.5 | 2.47 | 65.4 |  | Hutchings and Jones (1998) |
| Norway | Ogna | 8.8 | 13.2 | 2.65 | 54.6 |  | Hutchings and Jones (1998) |
| Norway | Orkla | 7.8 | 11.4 | 2.82 | 56.8 |  | Hutchings and Jones (1998) |
| Norway | Otra | 8.6 | 12.8 | 2.63 | 60.1 |  | Hutchings and Jones (1998) |
| Norway | Tengselva | 9.3 | 14.1 | 2.92 | 60.8 |  | Hutchings and Jones (1998) |
| Norway | Tovdalselva | 8.7 | 13 | 2.72 | 52.8 |  | Hutchings and Jones (1998) |
| Norway | Varhaugelva | 8.6 | 12.1 | 2.73 | 49.3 |  | Hutchings and Jones (1998) |
| Norway | Bouleau | 8.36 | 13.1 | 3.14 | 50.6 |  | Hutchings and Jones (1998) |
| Ireland | Burrishoole |  | 12.1 | 2.24 |  | 0.0375 | Bley and Moring (1988), Hutchings and Jones (1998) |
| Ireland | Corrib |  | 13.4 | 1.75 |  | 0.0266 | Hutchings and Jones (1998) |
| Scotland | North Esk |  | 12.6 | 2.33 |  | 0.0601 | Hutchings and Jones (1998) |

In addition, from Hutchings and Jones (1998), we selected data of populations with annual survival rates of smolt-grilse below 0.1 to ensure survival is a critical factor in the dynamics of those populations. Given that data on smolt-grilse survival is only available for a small number of populations (11 populations) and that for these populations the data on smolt age, length of 1+ parr and smolt is incomplete, we use only the smolt age and smolt length (if available) as a proxy of growth rate of presmolts. Smolt age is an approximate measure of the growth rate of presmolts, so the higher the smolt age the lower the growth rate as presmolts (Hutchings & Jones, 1998; Metcalfe & Thorpe, 1990; Power, 1981). The growth rate was calculated as the ratio between the smolt length and the smolt age. For those populations without information on smolt length (2 populations), we use the smolt length average of all populations (14.8 cm (Hutchings & Jones, 1998)). We tested the correlation between the growth rate of presmolts and the survival rate of postsmolts with a Pearson correlation test.

## Results

Under most conditions studied, the population exhibits a regular annual cycle (annual fixed point dynamics), meaning that the same values of population abundance and biomass, as well as food density in the breeding habitat, occur every year at a particular point of the year cycle (i.e. start of the year, spawning day, hatching day), even though within the year these densities, of course, vary. The population also exhibits 4 year cycles when conditions for postsmolts are very unfavorable, meaning a particular value of population abundance and biomass, and food density in the breeding habitat repeats itself every 4 years.

### Unfavorable conditions for postsmolts increase presmolt growth rate and affect life history trajectories

When the food in the non-breeding habitat is abundant (high feeding levels), the survival in the non-breeding habitat is high and the cost of the breeding travel is low, the offspring production by the population is high. Favorable conditions for postsmolts results in high offspring production due to the combination of 1) high growth rate of postsmolts which reach a large body size and hence a high fecundity (Fig. 2a), 2) long lifespan that increases the opportunities to reproduce (Fig. 2c) and 3) low energy expenditure during the breeding travel resulting in more energy reserves available for reproduction at arrival in the spawning grounds (Fig. 2e). High offspring production means a large number of presmolts competing for food, which causes a decline in the food abundance in the non-breeding habitat and therefore a low growth rate during the presmolt stage (Fig. 2b 2d 2f).

**Figure 2.**
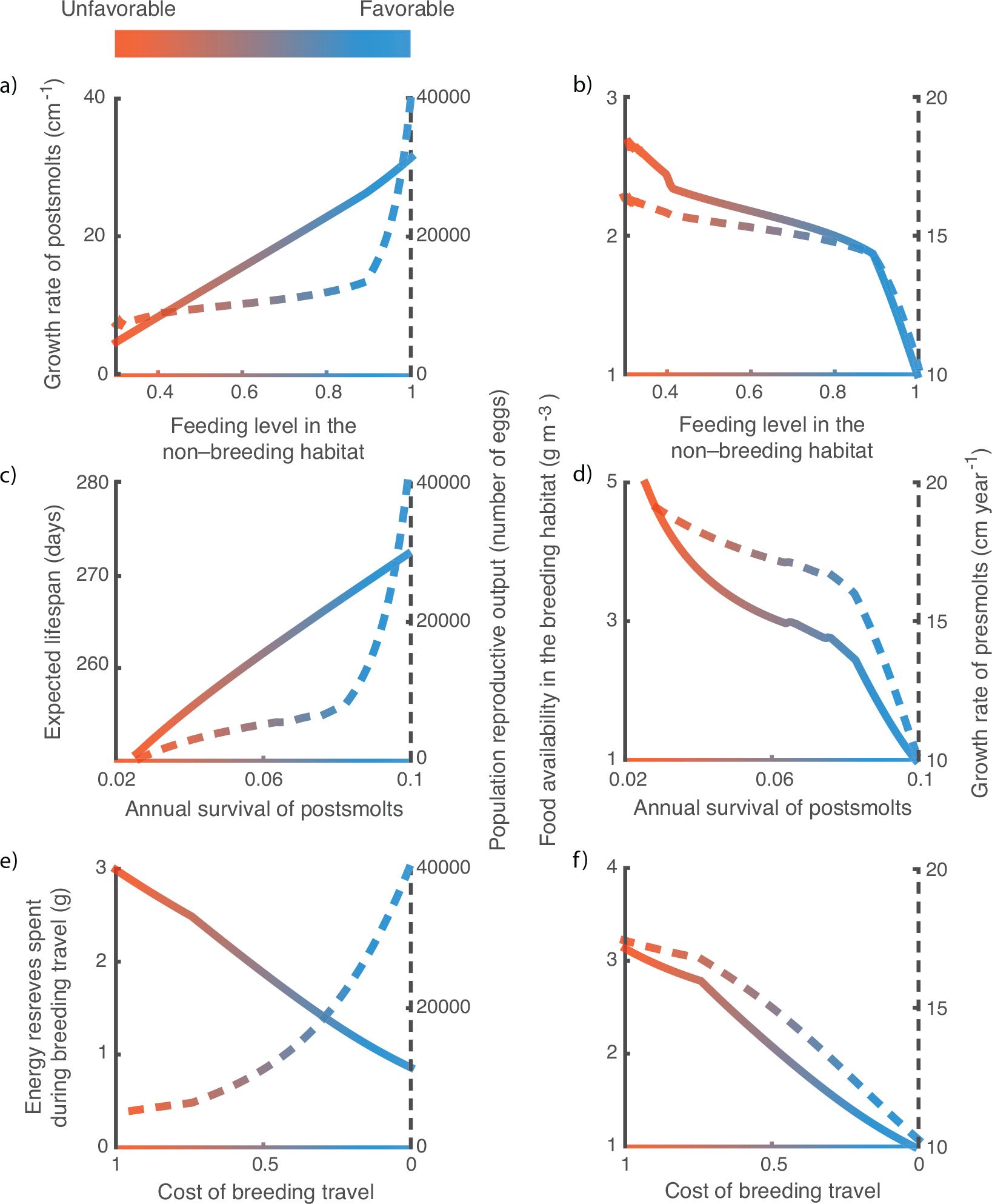
Effects of variation in feeding level in the non-breeding habitat (top row), annual survival of postmolts (middle row) and cost of the breeding travel (bottom row) on life history traits of postsmolts (solid lines in left column plots), population reproductive output (dashed lines in left column plots), food availability in the breeding habitat (solid lines in right column plots) and growth rate of presmolts (dashed lines in right column plots). Default values representing favorable conditions (feeding level in the non-breeding habitat = 1, annual survival of postsmolts = 0.1 and cost of the breeding travel = 0) are used for parameters that are not varied. *R*_*max*_= 5 g m^−3^, other parameter values as in table 2. The values correspond to the average population statistics after the transient dynamics have disappeared.

In contrast, low food abundance in the non-breeding habitat (low feeding levels) results in a decrease in the growth rate of postsmolts, which in turn causes a reduction in the population offspring production (Fig. 2a). Similarly, low survival of postsmolts reduces the individual lifespan and therefore the opportunities to reproduce, causing a low offspring production (Fig. 2c). Likewise, high cost of the breeding travel causes the individuals to spent a larger amount of their energy reserves during the migration, leaving less energy available for reproduction upon their arrival on the spawning grounds. Consequently, it results in low offspring production, and therefore in low densities of presmolts (Fig. 2e). Since a low number of presmolts means less competition in the breeding habitat, the food availability in this habitat is higher and therefore presmolts grow at higher rate (Fig. 2b 2d 2f) (Fig. 4).

Interestingly, despite that the three unfavorable conditions have the same consequence for presmolts, i.e. result in a high presmolt growth rate, their impacts on the life history trajectories of postsmolts differ. While low food abundance and low survival in the non-breeding habitat result in younger spawners (Fig. 3a 3b), high cost of the breeding travel does not affect the age at spawning (Fig. 3c). In the former case, the lower age at first spawning is a direct consequence of a high growth rate in the non-breeding habitat, which enables individuals to reach the maturation size faster and therefore to reproduce earlier. However, when the cost of the breeding travel is high, although the individuals after 1 year in the sea have reached the maturation size, the energy reserves they have accumulated after maturation only cover the high costs of the breeding travel, leaving no energy for offspring production. Effectively, they therefore postpone spawning until the reproductive season in the following year.

**Figure 3.**
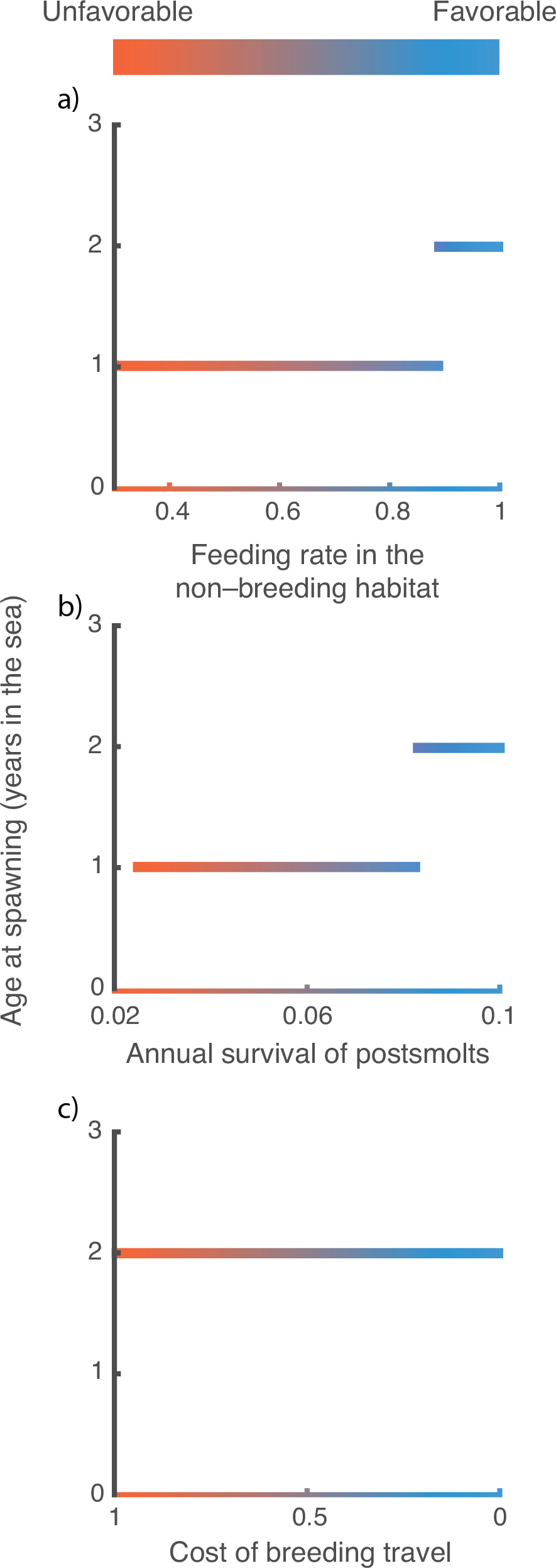
Effects of variation in feeding level in the non-breeding habitat (top row), annual survival of postmolts (middle row) and cost of the breeding travel (bottom row) on sea-age at first spawning. Same parameters as in figure 2. The values correspond to the average population statistics after the transient dynamics have disappeared.

### Unfavorable conditions for postsmolts affect population dynamics

Unfavorable conditions for postsmolts have effects on the population by affecting its abundance and its dynamics. For instance, a decrease in food abundance in the non-breeding habitat not only causes a decrease in the total population biomass, but also leads to the occurrence of population cycles with a period of 4 years when feeding levels are low (feeding level below 0.5 in Fig. 4a 4b). Similarly, low survival of postsmolts or high cost of the breeding travel also lead to the occurrence of such cycles (Fig. 4c 4d). These 4-years cycles are caused by the alternation of a fast and a slow-growing presmolt cohort. As mentioned above, unfavorable conditions for postsmolts result in low offspring production, which is very low when the conditions are extremely unfavorable, such as very low feeding levels in the non-breeding habitat. Very low offspring numbers result in high growth rates of presmolts due to low competition in the breeding habitat. Those individuals can reach large sizes before smolting and therefore accumulate enough energy that enables a high fecundity. This fast-growing cohort, therefore, produces a large number of offspring experiencing high competition in the breeding habitat, which causes the new cohort to grow slowly and accumulate less energy. As a consequence, this slow-growing cohort has low fecundity and produces an offspring cohort of small number that experiences low competition in the non-breeding habitat, resulting in a fast-growing cohort. The period cycle equals twice the age at first spawn (2 years).

**Figure 4.**
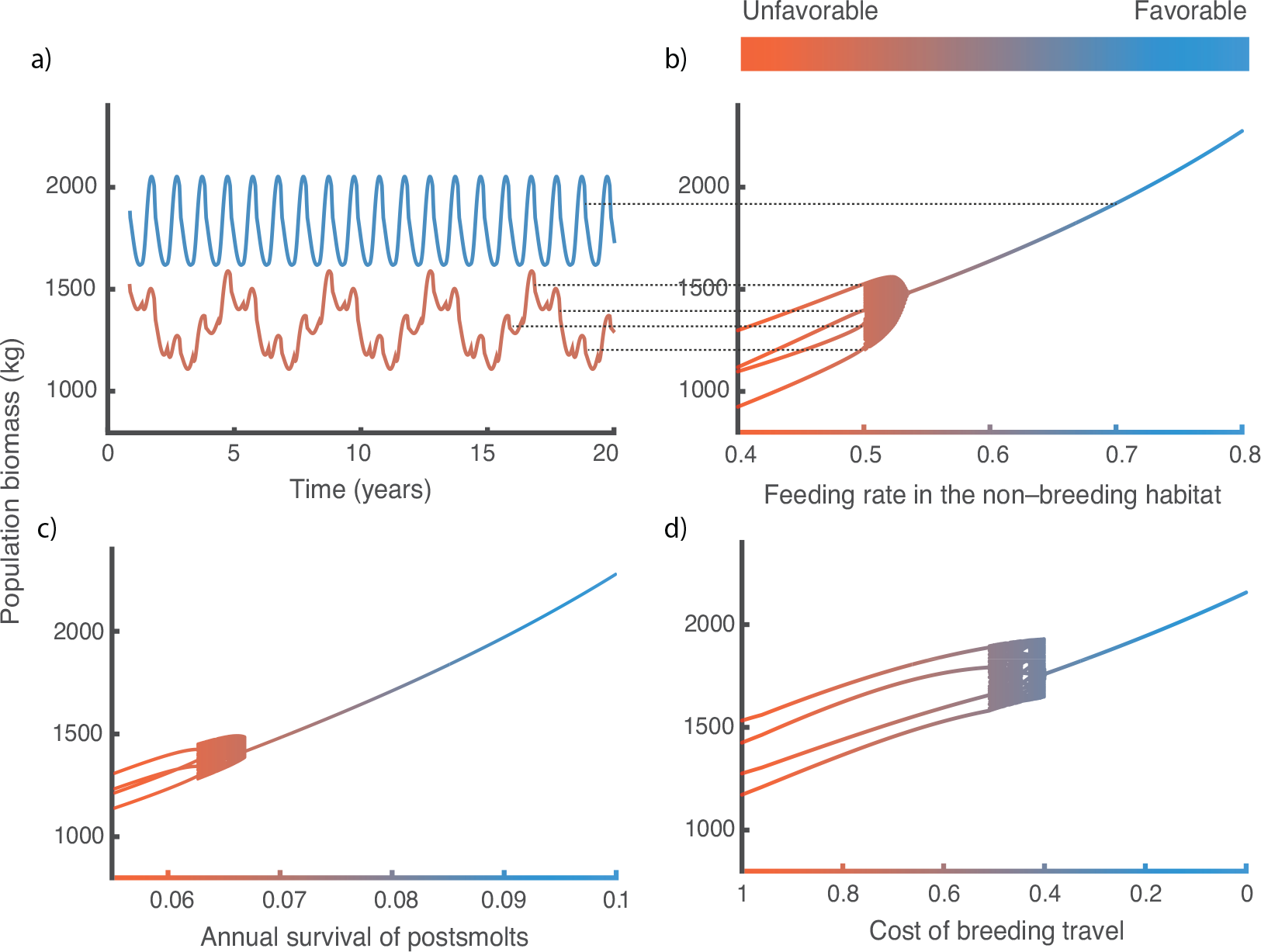
a), b) Effect of feeding level in the non-breeding habitat, c) annual survival of postmolts and d) cost of the breeding travel on population biomass dynamics. a) Annual fixed-point dynamics (blue) and 4 year cycles (brown) occur when feeding level in the non-breeding habitat equals 0.7 and 0.5 respectively. Dotted lines in a) indicate the time points in the dynamics at which the yearly census the population biomass occurs, resulting in the values shown in b). Default values representing favorable conditions (feeding level in the non-breeding habitat = 1, annual survival of postsmolts = 0.1 and cost of the breeding travel = 0) are used for parameters that are not varied. *R*_*max*_= 8 g m^−3^, other parameter values as in table 2. The values in plots b), c), and d) correspond to the population biomass census occurring every year at the time of hatching after the transient dynamics have disappeared.

Extremely unfavorable conditions for postsmolts are not the only cause of a shift in the dynamics from stable annual cycles to 4-years cycles, favorable conditions in conjunction with high maximum food density (carrying capacity) in the breeding habitat also cause the occurrence of oscillating dynamics (See SuppInfo 2). Since higher maximum food density in the breeding habitat results in higher growth potential of presmolts, the alternation of a weak and a strong cohort arises when conditions for postsmolts are not extremely unfavorable.

### Data from wild populations

Unfavorable conditions for postsmolts including low survival of postsmolts and low food abundance in the non–breeding habitat are correlated with high growth rates of presmolts. The growth rate of postsmolts (taken as a proxy for the food availability in the non–breeding habitat) is negatively correlated to the growth rate of presmolts (p<0.05, R-squared=−0.42) (Fig. 5a). Thus, when the food available in the non–breeding habitat is high and consequently the growth rate of postsmolts is high, the growth rate of presmolts is low. Conversely, a high growth rate of presmolts occurs when the growth rate of postsmolts is low. Similarly, the survival rate of postsmolts is negatively correlated to the growth rate of presmolts (p<0.05, R-squared=−0.61) (Fig. 5b), therefore an increase in survival of postsmolts decreases the growth rate of presmolts.

**Figure 5.**
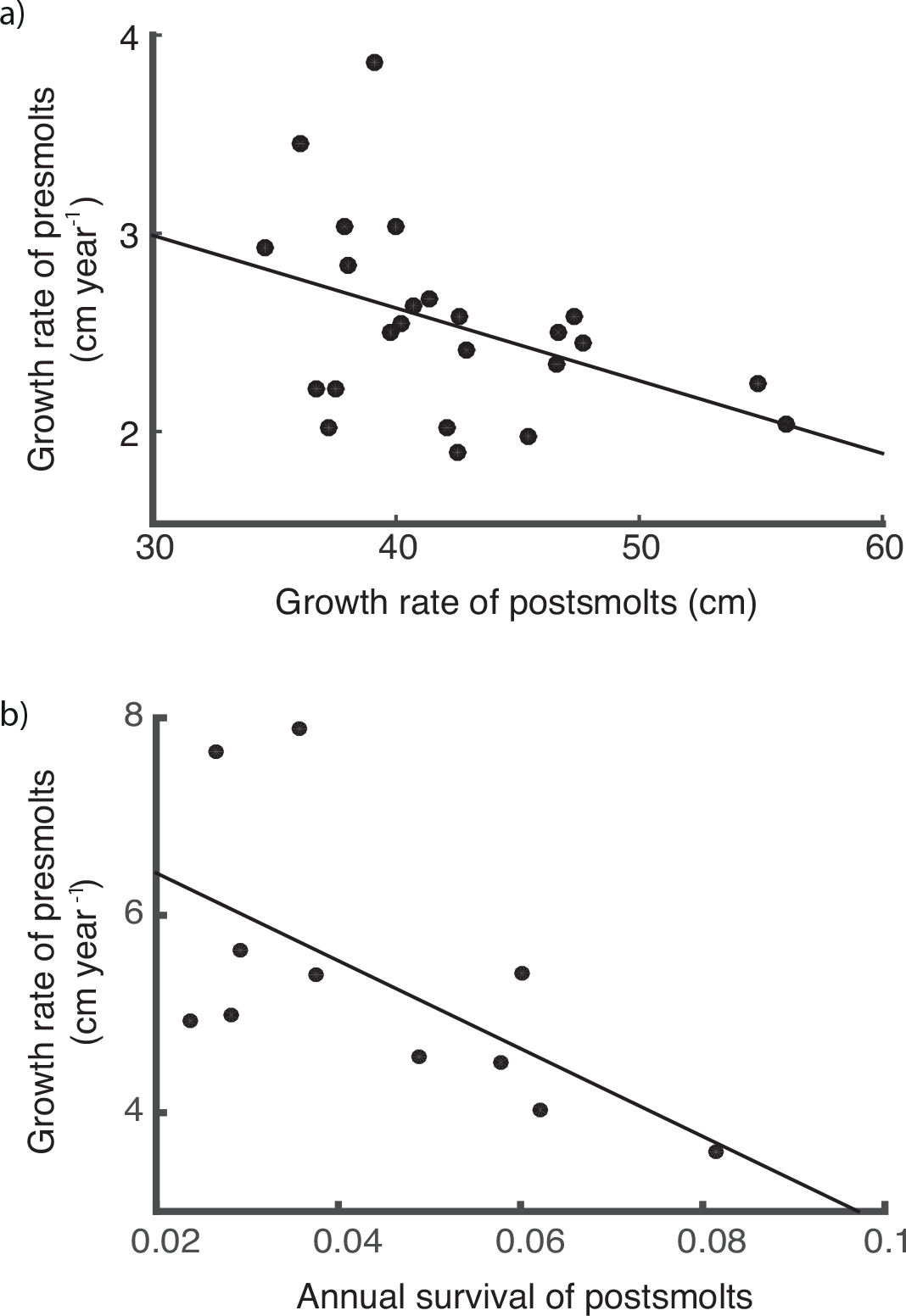
a) Growth rate of presmolts vs growth rate of postsmolts of 24 populations (Regression: p<0.05, R-squared=−0.42). b) Growth rate of presmolts vs annual survival of postsmolts of 11 wild populations (Regression: p<0.05, R-squared=−0.61).

## Discussion

Although relations between postsmolt conditions and presmolt growth rates in salmonids have rarely been studied in wild populations, the negative correlation between postsmolts and presmolts growth rates was previously described in a wild population (Einum, Thorstad, & Naesje, 2002). Similarly, the growth rate of postsmolts from the Drammen River have decreased in the last decades while the growth rate of presmolts have increased (McCarthy, Friedland, & Hansen, 2008). Yet, the mechanism that causes this difference in growth rates between different stages was not clear and different explanatory hypotheses have been proposed: the presence of traits having opposing effects on growth in different environments population (Einum et al., 2002) or a different growth rate in each environment due to a different necessity of escapement from different predation risk (Abrahams & Sutterlin, 1999). Here, we describe a simple mechanism arising from the interaction between individuals, in particular, the intimate link between the strength of density-dependence in the breeding habitat and the reproductive output of postsmolts (Fig. 6). Furthermore, given that the strength of the density-dependence in the breeding area determines the presmolt growth rate while the postsmolts growth rate directly affects the reproductive output, the presmolt and postsmolts growth rates are negatively correlated. Moreover, farmed Atlantic salmon that are not limited by food grow faster as presmolts compared to wild salmon (McGinnity et al., 2003), whereas both farmed and wild Atlantic salmon have similar growth rates in the ocean (Jonsson et al., 2003). Therefore, the difference in growth rates of individuals in wild populations in the two habitats is not an individual trait but is more likely arising from the interaction between individuals. This supports our hypothesis that density-dependence in the breeding habitat is the main cause of the negative correlation in growth rates rather than individual traits or predation risk.

**Figure 6.**
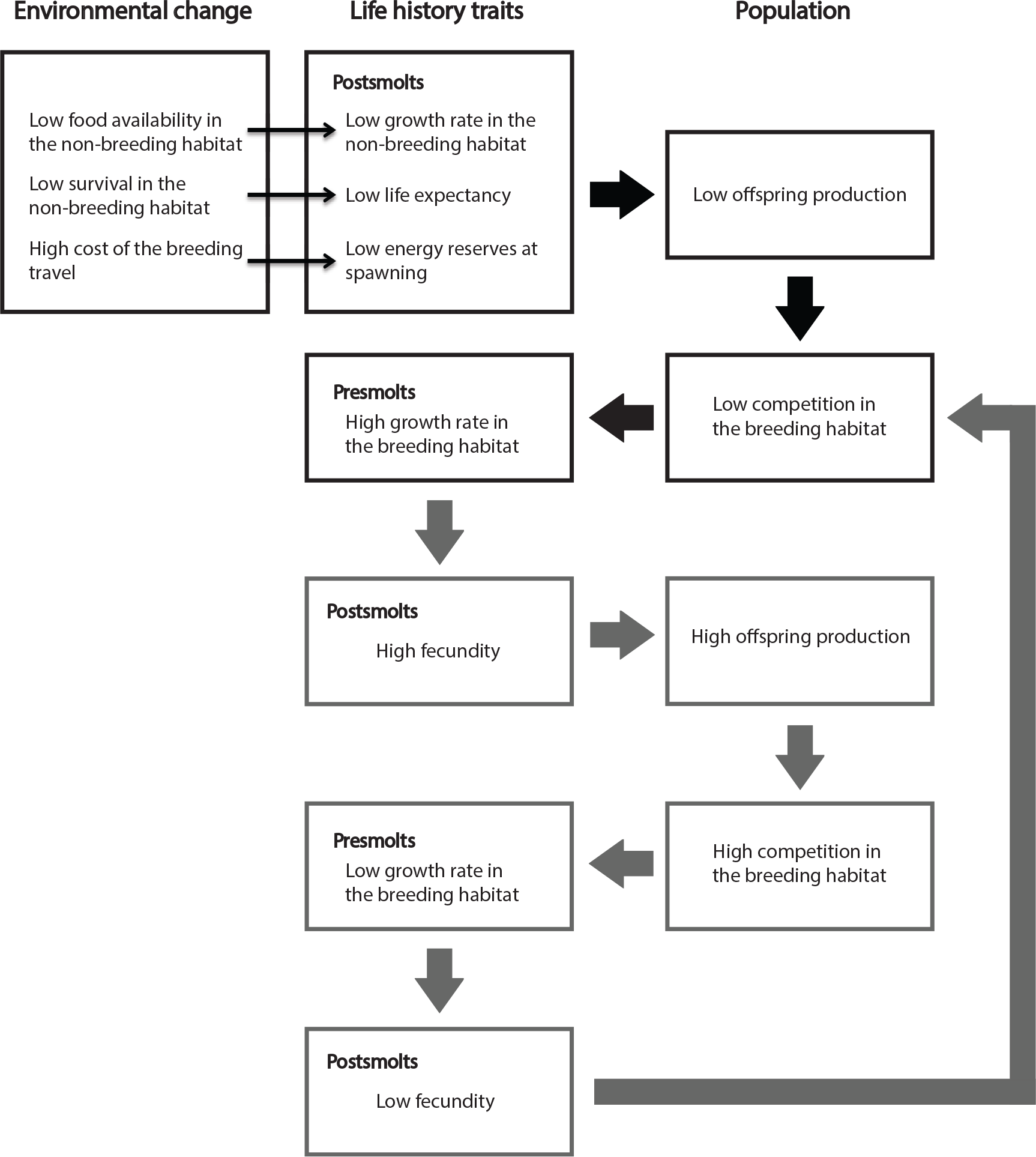
Direct (narrow arrows) and indirect (thick arrows) effects of environmental change in life history traits and population processes. Life history traits and population processes causing 4-years cycles in grey arrows and boxes.

Likewise, based on data of Atlantic salmon presented by Hutchings and Jones (1998), we show that low survival of postsmolts is correlated with high growth rate of presmolts as well, which is also predicted by our model as a consequence of reduced competition in the breeding habitat. These two unfavorable conditions also have an impact on the life history trajectory of individuals by reducing the sea-age at first spawning from 2 to 1 year. In fact, the average sea-age at first spawn has decreased in recent decades and this decrease coincides with a decline in the growth rate of postsmolts (Jonsson & Jonsson, 2004).

The effects of environmental change on wild populations, both in abundance (Lane, Kruuk, Charmantier, Murie, & Dobson, 2012; Ozgul et al., 2010) and in dynamics (Cornulier et al., 2013; Nelson & Yamanaka, 2013) have become increasingly documented. However, most analyses are phenomenological. Here, we focus on the mechanisms leading to changes in population abundance and dynamics of anadromous fishes facing environmental change through variation in food abundance, survival and cost of the breeding travel. In particular, we predict a change of dynamics from stable annual cycles to 4-years cycles in the face of unfavorable conditions for postsmolts or favorable conditions in combination with high food carrying capacity in the breeding habitat. Interestingly, cyclic dynamics have been observed in wild populations of other salmonid species. For instance, four year oscillations in some populations of Sockeye salmon (*Oncorhynchus nerka*) have motivated theoretical studies that explained the occurrence of this dynamics to be caused by tritrophic interactions (Guill, Drossel, Just, & Carmack, 2011), stochastic processes (Myers, Mertz, Bridson, & Bradford, 1998), depensatory predation or genetic effects (Levy & Wood, 1992). Our model successfully predicts the oscillations as a result of the density-dependent feedback between individuals within a cohort through the food density in the breeding habitat that causes the alternation of a fast and a slow-growing cohort (Fig. 6). The occurrence of 4-years cycles in populations of sockeye salmon coincides with a breeding habitat with high food carrying capacity rather than unfavorable conditions of postsmolts, according to empirical evidence and in line with previous theoretical studies (Guill et al., 2011). To our knowledge, there are no reported examples of transition of dynamics in anadromous populations to date. However, under a scenario of declining ocean productivity due to climate change (Hoegh-Guldberg & Bruno, 2010), we may observe in the future a destabilization of populations into cyclical dynamics. Such changes in population dynamics potentially will have consequences for the entire marine community the salmon is part of (Millon et al., 2014).

Simultaneous changes in life history traits and population dynamics in response to environmental change are common in nature (Parmesan, 2006). However, the causes underlying these joint responses remain largely unidentified. Few studies have shown the causal relation between coupled responses of individual traits and population dynamics to environmental change (Ozgul et al., 2010; Thompson & Ollason, 2001). In this study, we demonstrate how changes in life history traits and population dynamics are intimately linked. In particular, we show how environmental change, by affecting individuals in a specific life stage, has multiple consequences for other life stages and for the entire population. This is particularly relevant for species with complex life cycles involving more than one habitat; because they are exposed to different impacts of environmental change across the habitats and, therefore, across their life cycle. In this context, we provide a mechanistic understanding of the interactions between life history and population dynamics based on individual energy budgets for anadromous fishes. If we hope to accurately predict the biological consequences of environmental change, this mechanistic understanding is certainly required.

## Supporting information

Detailed population model and supplementary figure

## Acknowledgements

This research was supported by funding from the European Research Council (ERC) under the European Union’s Seventh Framework Programme (FP/2007-2013)/ ERC Grant Agreement no. 322814.

## Authors’ contributions

Chaparro-Pedraza and de Roos conceived the ideas, designed methodology, contributed critically to the drafts and gave final approval for publication; Chaparro-Pedraza analyzed the results and led the writing of the manuscript.

