## Supplementary material for "Environmental change effects on life history traits and population dynamics of anadromous fishes": Detailed population model and supplementary figure

**Supporting information 1. Physiologically–structured population model**

The physiologically–structured population model follows the cohort-based approach for populations with seasonal reproduction introduced by (Persson et al., 1998). Since reproduction occurs as a discrete event at a specific time in the year, all individuals that are born in the same reproductive event are equal. They are collected into a single cohort and assumed to grow at the same rate. Thus, we can describe the dynamics of each cohort $i \in \mathbb{N}$ by using a system of ordinary differential equations, which keeps track of the density of individuals $N_i$, their age $A_i$, their structural mass $W_i$ and their energy reserves storage $S_i$. Juveniles are defined as individuals with structural mass smaller than the structural mass at maturity $W_p$ and adults as individuals with structural mass equal or larger than $W_p$. For each cohort $i$, age is a monotonically increasing function of time,

$$\frac{d}{dt}A_i = 1 \qquad (1)$$

The age of the individuals determines the stage, which in turn, determines the differential equations that describe the variation in density of individuals, their structural mass and stored energy reserves. Equations (2), (3) and (4) define the dynamics of eggs, presmolts and postsmolts, respectively. The density of individuals decreases due to a mortality rate specific to each stage. In addition, the presmolts and postsmolts may die due to starvation. During the egg stage the structural mass and storage does not change. The dynamics of the structural mass and energy reserves storage in presmolts and postsmolts depend on the amount of food they encounter as well as the breeding travel period if they are adults.

for $0 \leq A_i < a_h$

$$\begin{cases} \frac{d}{dt}N_i = -\mu_e N_i \\ \frac{d}{dt}W_i = 0 \\ \frac{d}{dt}S_i = 0 \end{cases}$$

(2)

for $a_h \le A_i < a_s$

$$
\begin{cases}
\dfrac{d}{dt}N_i = \begin{cases} -\mu_r N_i & if\ \dfrac{S_i}{W_i} \ge q_S \\ -\left(\mu_r N_i + \varphi\left(q_S \dfrac{W_i}{S_i} - 1\right)\right) & if\ S_i > 0\ and\ \dfrac{S_i}{W_i} < q_S \\ -\infty & otherwise \end{cases} \\
\dfrac{d}{dt}W_i = \begin{cases} \zeta_W\left(\kappa\ \dfrac{R_r}{K + R_r}\ j_a\ {W_i}^{2/3} - j_m\ W_i\right) & if\ \kappa\ \dfrac{R_r}{K + R_r}\ j_a\ {W_i}^{2/3} > j_m\ W_i \\ 0 & otherwise \end{cases} \\
\dfrac{d}{dt}S_i = \begin{cases} (1-\kappa)\dfrac{R_r}{K + R_r}\ j_a\ {W_i}^{2/3} & if\ \kappa\ \dfrac{R_r}{K + R_r}\ j_a\ {W_i}^{2/3} > j_m\ W_i \\ \dfrac{R_r}{K + R_r} j_a\ {W_i}^{2/3} - j_m\ W_i & otherwise \end{cases}
\end{cases}
$$

(3)

for $a_s \le A_i$

$$
\begin{cases}
\dfrac{d}{dt}N_i = \begin{cases} -\mu_s N_i & if\ \dfrac{S_i}{W_i} \ge q_S \\ -\left(\mu_s N_i + \varphi\left(q_S \dfrac{W_i}{S_i} - 1\right)\right) & if\ S_i > 0\ and\ \dfrac{S_i}{W_i} < q_S \\ -\infty & otherwise \end{cases} \\
\dfrac{d}{dt}W_i = \begin{cases} \zeta_W\left(\kappa\ f_s\, j_a\ {W_i}^{2/3} - j_m\ W_i\right) & if\ c1\ and\ (\sim c2\ or\ \sim c3) \\ 0 & otherwise \end{cases} \\
\dfrac{d}{dt}S_i = \begin{cases} (1-\kappa) f_s\, j_a\ {W_i}^{2/3} & if\ c1\ and\ (\sim c2\ or\ \sim c3) \\ f_s\, j_a\ {W_i}^{2/3} - j_m\, W_i & if\ \sim c1\ and\ (\sim c2\ or\ \sim c3) \\ -(j_m\ W_i + C\ j_m\ W_i) & otherwise \end{cases}
\end{cases}
$$

(4)

In this last equation $c1$, $c2$ and $c3$ are the conditions $\kappa\ f_s\, j_a\ {W_i}^{2/3} > j_m\ W_i$, $t_{um} \le t \le t_{dm}$, and $W_p \le W_i$, respectively, while $\sim c1$, $\sim c2$ and $\sim c3$ refer to the situation that these conditions do not hold. When the conditions are true, the amount of assimilates necessary to meet metabolic maintenance from the $\kappa$ fraction are enough ($c1$), the current time corresponds to the breeding travel period ($c2$) and the cohort is adult ($c3$).

Whenever a juvenile cohort reaches the maturation size $W_i = W_p$, at a particular time $t = t_p$, a maturation event occurs. At a maturation event, the juvenile cohort becomes an adult cohort. This does not affect any cohort statistics:

$$\begin{cases} A_i(t_p) = A_i(t_p{}^-) \\ N_i(t_p) = N_i(t_p{}^-) \\ W_i(t_p) = W_i(t_p{}^-) \\ S_i(t_p) = S_i(t_p{}^-) \end{cases} \tag{5}$$

Reproduction occurs instantaneously at $t = n\, t_y + t_r$, where $n \in \mathbb{N}$. At a reproductive event, a new cohort is formed from the storage biomass of adults, if their storage exceeds the storage level corresponding to a storage:structural mass ratio equal to the storage:structural mass ratio at which the adults matured:

$$\begin{cases} A_0(t_{rn}) = 0 \\ N_0(t_{rn}) = \left( \sum\limits_{i \in \{j \le n | W_j \ge W_p\}} N_i \cdot \max\left( S_i - \frac{S_p}{W_p} W_i, 0 \right) \right) \frac{\zeta_e}{W_b} \\ W_o(t_{rn}) = \kappa\, W_b \\ S_0(t_{rn}) = (1 - \kappa)\, W_b \end{cases} \tag{6}$$

At the same time, all other cohorts are renumbered and the energy reserves storage of the adults is set to the amount that makes storage:structural mass ratio equal to the storage:structural mass ratio at maturity.

$$\begin{cases} A_{i+1}(t_{rn}) = A_i(t_{rn}{}^-) \\ N_{i+1}(t_{rn}) = N_i(t_{rn}{}^-) \\ W_{i+1}(t_{rn}) = W_i(t_{rn}{}^-) \\ S_{i+1}(t_{rn}) = \begin{cases} \min\left( S_i(t_{rn}{}^-), \frac{S_p}{W_p} W_i(t_{rn}{}^-) \right) & if\ W_i \ge W_p \\ S_i(t_{rn}{}^-) & otherwise \end{cases} \end{cases} \tag{7}$$

The resource density in the breeding habitat grows following a semi–chemostat growth and declines by foraging of presmolts (8).

$$\frac{d}{dt}R_r = \rho(R_{max} - R_r) - \frac{R_r}{K + R_r} j_a \sum_{i \in \{j \le n | a_h < a_j < a_s\}} N_i \, W_i^{2/3} \tag{8}$$

**Supporting information 2.**

Figure SI2.1. Dynamics (left column) and total biomass (right column) of a population exposed to different feeding levels in the non-breding habitat (top row), cost of the breeding travel (middle row) and survival of postsmolts (bottom row) and variation in maximum food density in the breeding habitat (horizontal axes). Default values representing favorable conditions (annual survival of postsmolts = 0.1 and cost of the breeding travel = 0) are used for parameters that are not varied. Feeding level equals 0.6 and 0.8 in middle and bottom plot rspectively.The values correspond to the average population statistics after the transient dynamics have disappeared.

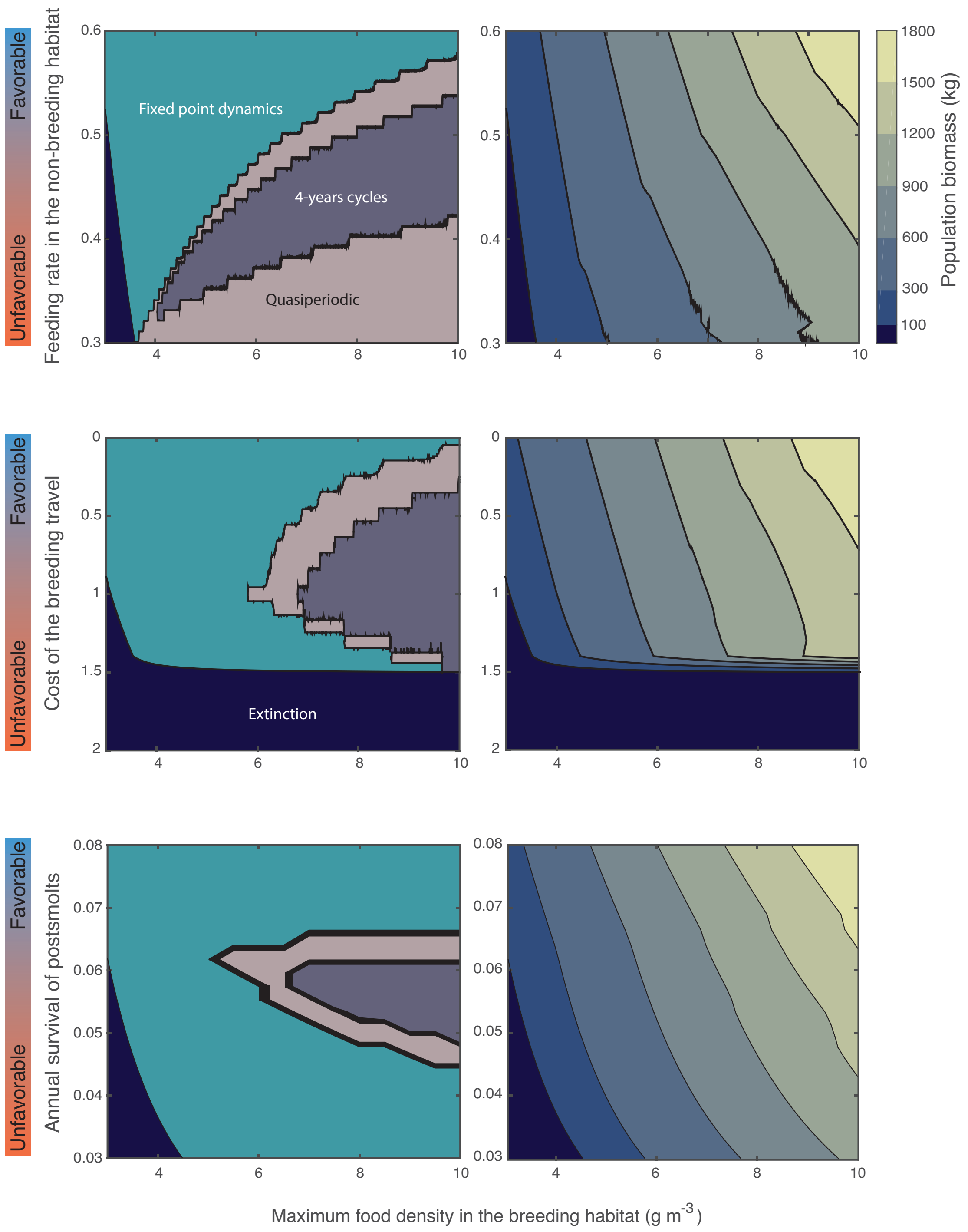
Favorable
Unfavorable
Feeding rate in the non-breeding habitat
Fixed point dynamics
4-years cycles
Quasiperiodic
Population biomass (kg)
1800
1500
1200
900
600
300
100
Cost of the breeding travel
Extinction
Annual survival of postsmolts
Maximum food density in the breeding habitat (g m-3)

59
